# Evaluating transcriptome-wide information preservation by ensilication using long-read RNA sequencing

**DOI:** 10.64898/2026.04.13.717197

**Authors:** Megha Muraleedharan, Harsha Gowda, James L. Banal

## Abstract

**Background:** Evaluation of RNA preservation requires an understanding of how storage conditions affect recoverable transcriptome information. Long-read RNA sequencing enables comprehensive assessment of transcript representation, isoform composition, and support for annotated transcript structures. This approach was employed to investigate the protective capabilities of ensilication during ambient and elevated-temperature storage.

**Results:** Pooled human total RNA was analyzed using PacBio Kinnex across six conditions, with two replicates per storage condition. Comparisons included frozen controls as well as ensilicated and unprotected RNA stored for seven days at room temperature or 40 °C. At matched input depth, with at least one qualifying sequence per transcript, ensilicated samples retained 88–89% of a fixed frozen-reference transcript set at room temperature and approximately 85% at 40 °C. In contrast, unprotected RNA retained 78–83% and 42–64%, respectively. Ensilicated samples exhibited smaller changes in isoform composition among evaluable genes. The retention advantage persisted when sequences were required to support a uniquely distinguishing complete intron chain and both annotated transcript boundaries. Retention estimates varied with detection criteria, and substantial limitations in annotated end-to-end coverage were observed even in frozen controls.

**Conclusion:** Long-read RNA sequencing demonstrates that ensilication limits storage-associated loss and distortion of recoverable transcriptome-wide information in the tested RNA preparation. These findings indicate a protective effect during seven days of ambient and elevated-temperature storage.

## Background

Long-read RNA sequencing characterizes full-length transcript isoforms from individual RNA or cDNA molecules [1–3]. Iso-Seq reads can span entire cDNA molecules, providing evidence for alternative splicing and transcript boundaries that are often difficult to reconstruct from short reads [4]. The LRGASP consortium demonstrated that longer, more accurate reads improve transcript identification, while greater sequencing depth enhances quantification [5]. RNA integrity is therefore critical when evaluating how preservation methods retain the information required for long-read transcriptome analysis.

RNA degradation affects both short-read and long-read measurements. Short-read protocols using random-primed cDNA synthesis can recover gene-level information from substantially degraded samples [6, 7]. In the Iso-Seq workflow, full-length reads are identified by recognition of library primers and the poly(A) sequence [8]. A cDNA molecule generated from a poly(A)-bearing RNA fragment can carry both library primers. Pipeline-defined full-length non-chimeric (FLNC) status must therefore be distinguished from coverage of an intact native transcript. Nevertheless, fragmentation can remove features needed to distinguish isoforms and alter their measured representation. A controlled degradation study using nanopore direct RNA sequencing demonstrated reduced library complexity and overrepresentation of shorter genes and isoforms [9]. Maintaining RNA quality during storage and transport is consequently an important practical consideration for transcriptomic studies. Storage at −80 °C is widely used to preserve RNA but requires continuous refrigeration and specialized equipment [10]. These requirements constrain decentralized sample collection, field research, and biobanking, motivating the evaluation of ambient-temperature preservation approaches.

Several approaches have been developed for ambient-temperature RNA preservation. RNAlater preserves RNA for days to weeks [11, 12], although studies of stored tissues have reported non-random changes in gene expression relative to frozen samples [13, 14]. Desiccation matrices such as RNAstable support prolonged room-temperature storage and have yielded gene-expression and short-read sequencing results comparable to frozen samples [10, 15, 16]. These studies demonstrate compatibility with downstream assays, but provide limited direct evidence for retention of complete transcript structures. Gene-level information can remain recoverable after fragmentation has removed features needed to distinguish individual isoforms. Long-read RNA sequencing therefore provides a complementary approach for evaluating how preservation affects transcript representation, isoform composition, and structural support across the transcriptome.

Ensilication encapsulates biomolecules within a deposited silica matrix, rendering them thermally stable within a conformal cage from which they are released intact [17–19]. For DNA, it maintains diagnostic concordance with cryogenic controls, accumulates fewer artifactual C-to-T mutations [20], and supports native long-read sequencing comparable to frozen across read length, fidelity, and methylation[21]. For RNA, prior silica-based preservation studies demonstrated recovery using RT-qPCR, Sanger sequencing, and targeted short-read amplicon sequencing [22–24]. These findings establish that RNA-derived sequence information can be recovered after encapsulation. However, they do not provide a transcriptome-wide assessment of how ensilication protects isoform representation, composition, and support for annotated transcript structures during storage.

Here, we used PacBio Kinnex long-read RNA sequencing to evaluate transcriptome-wide information preservation by ensilication. Pooled human total RNA was profiled across six conditions comprising frozen storage at −80 °C, an ensilication-and-release control, and ensilicated or unprotected RNA stored for seven days at room temperature or 40 °C. The primary preservation benchmark focused on annotated transcripts. We assessed gene and transcript representation, within-gene isoform composition, and sequence support for annotated intron chains and transcript boundaries. At matched input depth and the primary detection threshold, ensilicated samples retained more frozen-reference transcripts than unprotected samples and showed smaller isoform-composition changes among evaluable genes. These analyses characterize the protective capabilities of ensilication in terms of recoverable transcriptome-wide information under the tested storage conditions.

## Methods

### RNA source material

Commercially available pooled human total RNA (BioChain Universal Human RNA, catalog number: R4234565-1, lot: C811087) served as the input for all experimental conditions. This RNA was derived from major organs of 10 adult donors (6 male, ages 27–76; 4 female, ages 27–66), isolated using modified guanidine thiocyanate extraction, treated with DNase I, and supplied at a concentration of 1.00 µg/µL. A working stock was prepared at 100 ng/µL in 10 mM sodium citrate with 0.01% Tween-20. We prepared all experimental conditions in duplicate from the same working stock.

### Ensilication and storage

For each ensilicated sample, 10 µg of total RNA was combined with ensilication microparticles (Cache DNA, Inc.) and mixed by hand inversion. Ensilication reagent (Cache DNA, Inc.) was added and samples were vortexed at 1,400 rpm for 2 days at room temperature. The resulting particle suspensions were stored in water. For the ensilicated control condition, samples were stored for 1 h following encapsulation, then pelleted at 1,000 × g for 1 minute and processed for de-ensilication. The encapsulation schedule ensured that both the 1-hour control and the 7-day stored ensilicated samples were de-ensilicated on the same day. For room temperature storage, ensilicated suspensions were maintained at ambient conditions for 7 days before processing. For accelerated thermal stress, ensilicated suspensions were transferred to an environmental chamber (LH 1.5, Associated Environmental Systems) and stored at 40 °C and 70% relative humidity for 7 days before processing. Requests for access to ensilication reagents should be directed to Cache DNA, Inc.

### De-ensilication and recovery

Following storage, particles were resuspended in 100 µL of de-ensilication buffer. The resulting solution was desalted twice using 4 mL 10 kDa MWCO spin filters (Amicon Ultra-4) with 10 mM sodium citrate and 0.01% Tween-20 as the wash buffer. The recovered RNA was used directly for library preparation. Requests for de-ensilication reagents should be directed to Cache DNA, Inc.

### Bare RNA storage

For bare RNA conditions, 10 µg of total RNA in 10 mM sodium citrate and 0.01% Tween-20 was aliquoted into 1.5 mL microcentrifuge tubes without preservative. Room-temperature samples were stored under ambient conditions for 7 days. Accelerated thermal stress samples were stored in an environmental chamber (LH 1.5, Associated Environmental Systems) at 40 °C and 70% relative humidity for 7 days. Frozen reference samples were stored at -80 °C. All samples were used directly for library preparation following storage.

### RNA quality assessment

RNA integrity was assessed using a 4200 TapeStation system (Agilent) with High Sensitivity RNA ScreenTapes. RINe values were calculated using the TapeStation Analysis Software. RNA recovery and purity metrics are reported in Supplementary Table 1.

### Library preparation and sequencing

Full-length cDNA was generated from total RNA input using the PacBio Iso-Seq Express 2.0 kit with oligo-dT priming. Sequencing libraries were prepared using the PacBio Kinnex full-length RNA kit according to the manufacturer’s protocol. Library preparation and sequencing were performed by Signios Biosciences (Foster City, CA), with all twelve sample libraries prepared concurrently. The two barcoded replicates from each condition were pooled for sequencing across six SMRT Cells, with each condition assigned to a separate sequencing pool.

### Data processing and analysis

Raw HiFi reads were segmented into individual transcript reads using Skera v1.2.0 (https://github.com/PacificBiosciences/skera). Segmented reads underwent primer identification and removal with lima v2.10.0 (https://github.com/PacificBiosciences/barcoding) using the --isoseq option, followed by Iso-Seq v4.1.2 refinement with --require-polya to obtain full-length non-chimeric (FLNC) reads. FLNC reads were clustered into transcript consensus sequences using isoseq cluster2 (Iso-Seq v4.0.0; https://github.com/PacificBiosciences/IsoSeq). Clustered reads were aligned to GRCh38 (GCA_000001405.15, no alt analysis set; https://ftp.ncbi.nlm.nih.gov/genomes/all/GCA/000/001/405/GCA_000001405.15_GRCh38/) using pbmm2 v1.14.99 (https://github.com/PacificBiosciences/pbmm2) with the --preset ISOSEQ --sort flags and collapsed into non-redundant isoform models using isoseq collapse.

Gene-level and transcript-level quantification was performed using IsoQuant v3.5.1 (https://github.com/ablab/IsoQuant)[25] with the --data_type pacbio_ccs and --complete_genedb flags against the GENCODE v43 comprehensive annotation (https://www.gencodegenes.org/human/release_43.html)[26]. The primary preservation analyses used annotated reference transcripts and their assignment counts. Isoform structural classification was performed using SQANTI3 v6.0 (https://github.com/ConesaLab/SQANTI3)[27]. Unless otherwise specified, downstream sequence counts refer to clustered consensus sequences without weighting by their numbers of contributing FLNC member reads.

For Figure 1c, sequence lengths were obtained from the length field of the Iso-Seq collapse read-statistics files. Each row contributed the corresponding FLNC member read’s own post-refinement sequence length following primer and poly(A) processing. The plotted population therefore comprised pre-clustering FLNC reads represented in the collapse output. The two replicates from each condition were pooled without additional filtering or subsampling, and medians and interquartile ranges were calculated from the represented member-read lengths. Read-length summaries and comparisons were descriptive only, without statistical tests treating individual reads as independent preservation replicates.

**Figure 1.**
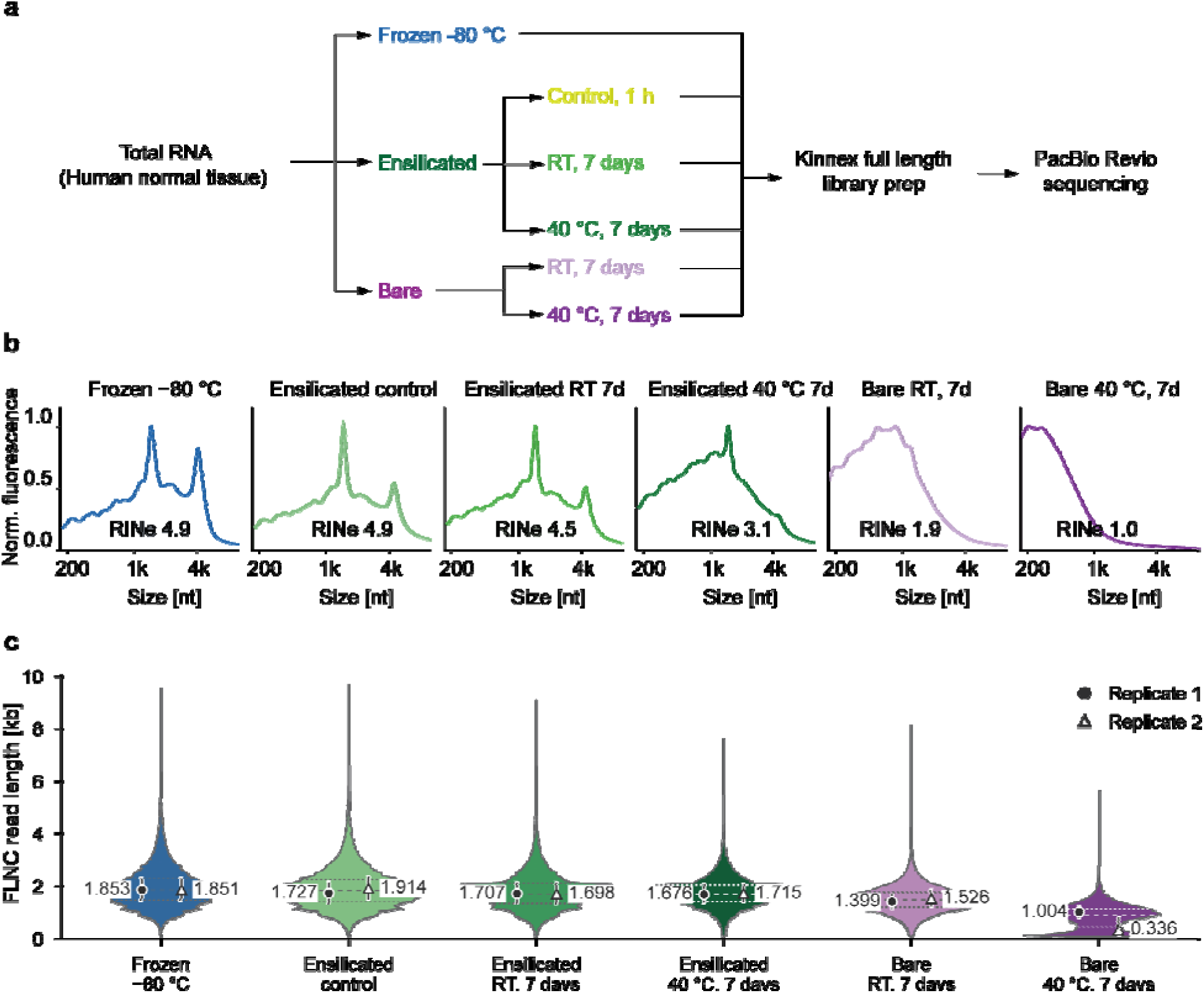
RNA integrity and FLNC sequence lengths following ensilication and storage. **a**, Experimental design. Pooled human total RNA was stored under six conditions in duplicate and processed through a common Kinnex full-length RNA library preparation and PacBio Revio sequencing workflow. **b**, TapeStation electropherograms display one representative sample per condition, with RINe values annotated in each panel. Electropherograms for both samples are provided in Supplementary Fig. 1. **c,** Length distributions of pre-clustering full-length non-chimeric (FLNC) reads are shown from the Iso-Seq collapse output. Each violin plot pools post-refinement member-read lengths from the two barcoded replicates per condition. Lengths are presented in kilobases. Dashed lines indicate the pooled 25th, 50th, and 75th percentiles. Filled circles and open triangles represent the medians for replicates 1 and 2, respectively, with vertical bars indicating the corresponding interquartile intervals. Numerical labels denote the individual sample medians.

### Gene and transcript abundance analysis

Gene abundance was expressed as counts per million gene-assigned consensus sequences (CPM). The denominator was the sum of counts across all genuine gene rows before expression filtering, excluding assignment and alignment summary rows. No transcript-length correction was applied. Principal component analysis was performed on log_2_(CPM + 1)-transformed gene abundances using singular value decomposition. Spearman rank correlations were calculated for pairwise sample comparisons at gene and annotated-transcript levels, retaining zero values within the common feature sets. The numbers of features included and the treatment of non-feature summary rows are reported in Supplementary Table 2.

### Annotated transcript retention

A fixed frozen-reference set was defined by requiring at least five annotated-transcript assignment counts and an abundance of at least 1 CPM in each frozen library. This selection yielded 13,362 transcripts. The same reference denominator was retained across samples, including transcripts with zero counts after storage. Count-only retention used the original IsoQuant annotated-transcript counts without additional structural-support filtering. Expected retention was evaluated at a common target input depth equal to the smallest consensus-sequence library, comprising 534,109 sequences. For each sample, the probabilities of meeting the detection threshold were summed across reference transcripts and expressed as a percentage of the fixed reference set.

Depth matching used exact hypergeometric detection probabilities under sampling without replacement. For each transcript, the retained count followed (*X* ∼ Hypergeometric(*N*, *c*, *n*)). The parameters were the sample’s observed input depth (*N*), the transcript’s qualifying count (*c*), and the target depth (*n*). Detection probability P_detect_ was P_detect_ = Pr(*X* >=*k*), where *k* was the required sequence count.

The primary detection threshold required at least one qualifying consensus sequence per transcript. Sensitivity analyses used thresholds of two, five, and ten sequences and the lower input depths shown in Supplementary Figure 6. Each library was evaluated separately. Transcript lengths were calculated from the summed lengths of annotated exons. For the adjusted length analysis, transcripts were jointly stratified by frozen starting-abundance quartile and the number of annotated isoforms per gene, grouped as 1, 2–5, 6–10, and more than 10. Retention within each length bin was standardized to the same joint-stratum distribution across the full frozen-reference set. Fixed stratum denominators retained transcripts with zero detection. The shortest length bin was omitted from the adjusted comparison because it lacked reference transcripts in several joint strata.

### Differential gene abundance

Gene abundance comparisons were performed in R v4.5.1 using limma v3.68.5 (https://github.com/bioc/limma)[28]. The model used log_2_(CPM + 1) values without additional normalization and included genes with CPM greater than 5 in at least one library. A model with six condition means, specified as ∼0 + condition, was fitted across all twelve libraries. Five contrasts compared the non-frozen conditions with frozen storage. Moderated t tests used empirical Bayes estimation with trend=TRUE and robust=TRUE, with all twelve libraries contributing to variance estimation. A single Benjamini–Hochberg adjustment was applied across all 99,415 tests arising from 19,883 genes and five contrasts. Libraries were the analysis units, and the model assumed independent library errors. Condition and sequencing-pool effects could not be estimated separately.

For each comparison, the displayed genes were restricted to those with CPM greater than 5 in at least one of the two condition libraries or two frozen libraries. Classification as higher or lower abundance required an adjusted P value below 0.05. The corresponding estimated log₂ contrast had to exceed 1 or fall below −1, respectively. Remaining displayed genes were classified as other eligible genes. For Figure 4b, each gene was assigned a fixed exonic length weighted by its annotated isoform abundances in the frozen libraries. Genes without an evaluable frozen-reference length were excluded from the length summaries but retained in the abundance analysis. Length distributions were summarized using box plots and medians, without additional tests comparing gene lengths between groups.

### Isoform composition and dominant-isoform discordance

The frozen-eligible gene set required at least 20 annotated-transcript assignment counts per gene in each frozen library, and at least two shared annotated transcript identifiers, each supported by at least two counts in both frozen libraries. These criteria yielded 2,729 genes. Within each gene, isoform proportions were calculated by dividing each annotated transcript’s count by the sum of annotated-transcript counts for that gene, retaining zero-count isoforms. For non-frozen samples, the reference was the equal-weight mean of the two frozen proportion vectors. Isoform-composition distance was calculated as D = (1/2) sum_i_ |*p*_i_ - *q*_i_|, where *p*_i_ and *q*_i_ are the sample and reference proportions of isoform (i). Genes with zero assigned counts in a sample had undefined composition distance and were reported separately as absent. The two frozen libraries were also compared directly to provide a reference for between-library variation.

The dominant isoform was defined as the annotated transcript with the highest within-gene proportion. Dominant-isoform discordance was calculated as the fraction of evaluable genes for which this identity differed between the sample and reference. Genes absent from the sample or tied for the highest proportion in either vector were excluded from the denominator. Ties were identified using an absolute tolerance of 10^-12^ and zero relative tolerance. Sensitivity analyses additionally required the difference between the two highest isoform proportions to exceed 0.1 or 0.2 in both compared vectors. The direct frozen comparison used its own evaluable gene set under each corresponding criterion.

### Annotated transcript coverage and structural support

Coverage analyses used consensus sequences uniquely assigned to annotated transcripts by IsoQuant, including the unique and unique_minor_difference assignment categories, for which alignment and annotation boundaries could be compared. Missing annotated exonic sequence was calculated separately at the 5’ and 3’ ends, accounting for transcript strand. These measurements summed annotated exonic bases beyond the corresponding alignment end and excluded intervening intronic sequence. Missing lengths were summarized in kilobases for each sample and described annotation-relative coverage among captured, assigned consensus sequences.

Structural support was evaluated using three nested criteria. Distinguishing-junction support required every observed junction to be consistent with the assigned annotation and the intersection of the annotated transcript sets sharing those junctions to contain only the assigned transcript. Complete-chain support additionally required the entire annotated intron chain. The strict criterion further required both original genomic alignment ends to lie within 50 nucleotides of the corresponding annotated genomic boundaries. Monoexon transcripts and transcripts unresolved by junction evidence could not satisfy these criteria. Strict transcript retention used the same frozen-reference denominator and common input-depth target as count-only retention. Sequence-level pass fractions in Supplementary Figure 7 instead used all input consensus sequences in each library at its observed depth as the denominator. Sampling variation conditional on the observed libraries was distinguished from variation between experimental preparations.

### Genome browser visualization

BigWig coverage tracks were generated from genome-aligned BAM files using deepTools bamCoverage v3.5.6 (https://github.com/deeptools/deepTools)[29] with CPM normalization and 50-base-pair bins. Coverage tracks from the two samples within each condition were averaged using deepTools bigwigAverage v3.5.6. Splice junctions were extracted from BAM CIGAR strings using pysam (https://github.com/pysam-developers/pysam), retaining junctions supported by at least three aligned sequences. Browser panels were generated using pyGenomeTracks v3.9 (https://github.com/deeptools/pyGenomeTracks)[30] with the GENCODE v43 annotation. Junction arcs summarize support for individual splice junctions.

## Results

### Experimental design and RNA integrity

To evaluate whether ensilication preserves RNA integrity for long-read isoform sequencing, commercially available pooled human total RNA from major organs of 10 adult donors (6 male, 4 female, ages 27–76) was analyzed. This pooled material provided transcriptomic diversity across tissues, facilitating assessment of transcript detection and quantitative representation. Two replicates per condition were examined across six storage conditions using PacBio Kinnex sequencing (Fig. 1a). Frozen samples stored at −80 °C served as the reference. Ensilicated RNA was stored for seven days at room temperature or 40 °C, with an additional ensilicated control stored for 1 hour after encapsulation. Bare RNA was stored for seven days at the same temperatures to evaluate changes occurring without ensilication.

The effect of storage conditions on RNA integrity was evaluated using capillary electrophoresis (Fig. 1b and Supplementary Fig. 1). The preservation comparison employed partially degraded RNA, with RINe values of 4.9–5.0 in the frozen reference samples. RINe values remained similar in the ensilicated control (4.8–4.9) and after seven days of ensilicated storage at room temperature (4.5–4.7). Storage at 40 °C led to a greater decline in the ensilicated samples, with RINe values of 3.1–3.6. At both temperatures, ensilicated RNA exhibited higher integrity than bare RNA, which showed RINe values of 1.8–1.9 at room temperature and 1.0–2.3 at 40 °C. These results demonstrate that ensilication reduces further degradation during storage.

To assess whether differences in RNA integrity correlated with changes in recovered sequence lengths, post-refinement FLNC member reads from the Iso-Seq collapse output were analyzed, with sample medians and interquartile intervals (Q1–Q3) reported separately (Fig. 1c). Frozen samples exhibited median lengths of 1.853 (1.456–2.279) and 1.851 (1.455–2.280) kb, compared with 1.727 (1.343–2.126) and 1.914 (1.507–2.339) kb in the ensilicated control. After seven days of ensilicated storage, the corresponding values were 1.707 (1.350–2.109) and 1.698 (1.332–2.109) kb at room temperature, and 1.676 (1.384–2.035) and 1.715 (1.416–2.057) kb at 40 °C. Bare RNA showed greater shortening, with median lengths of 1.399 (1.147–1.661) and 1.526 (1.242–1.831) kb at room temperature, and 1.004 (0.804–1.218) and 0.336 (0.149–0.775) kb at 40 °C. The shift toward shorter captured sequences was therefore more pronounced following bare storage, particularly at 40 °C.

### Gene-level expression is preserved across all ensilicated conditions

RNA degradation can alter measured gene abundances because transcripts differ in susceptibility [6]. To assess whether differences in RNA integrity and recovered sequence lengths were associated with changes in gene-level abundance profiles, principal component analysis of log2(CPM + 1)-transformed abundances was performed to summarize the overall differences among the twelve libraries (Fig. 2a). PC1 and PC2 explained 47.5% and 11.6% of the total variance, respectively. Frozen and all three ensilicated conditions clustered together, while bare samples stored at room temperature occupied an intermediate position along PC1 between this group and the bare 40 °C samples. The two bare 40 °C samples also separated substantially from each other, indicating greater variation between their measured gene-abundance profiles.

**Figure 2.**
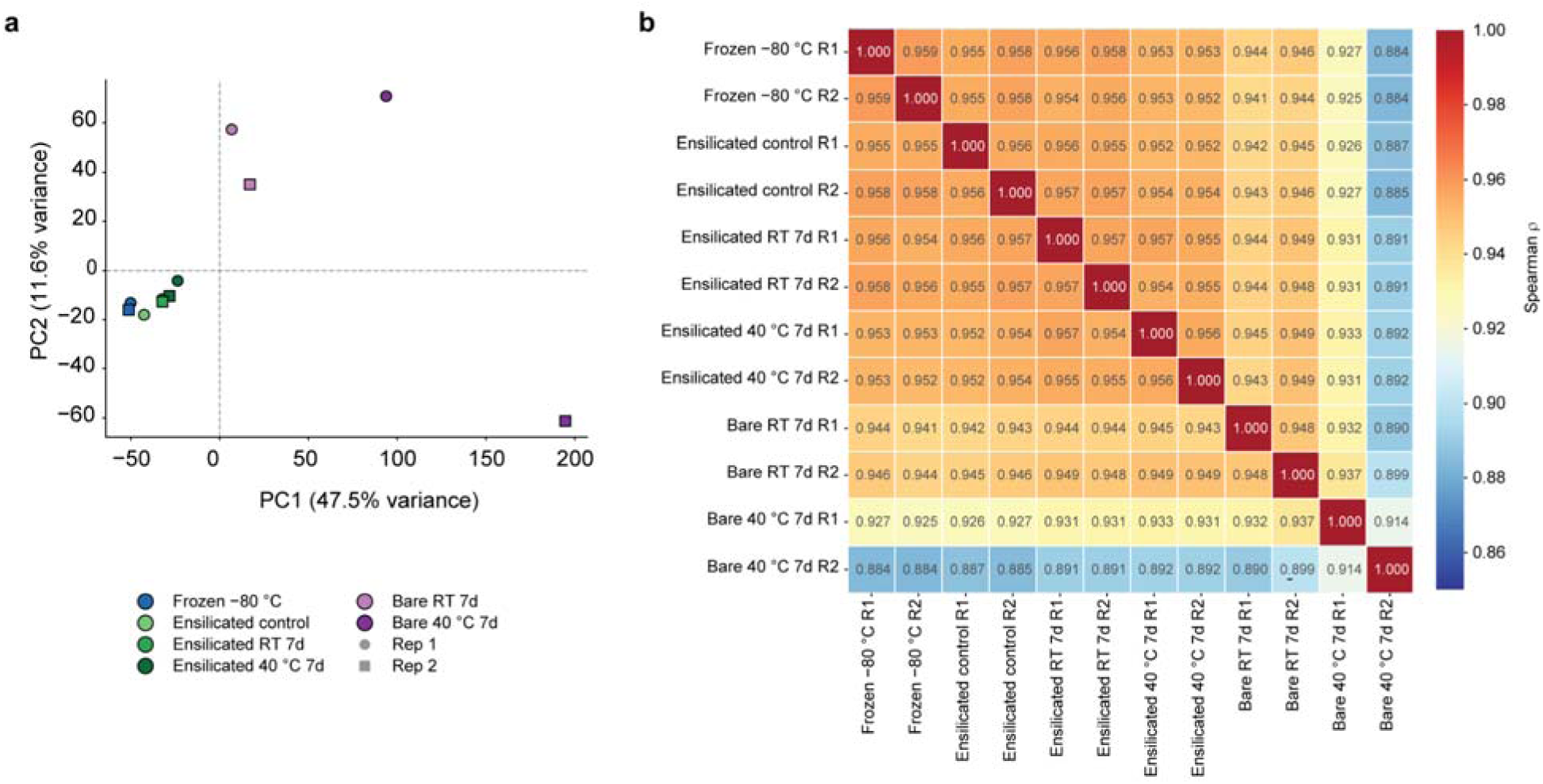
Gene-level abundance profiles following ensilication and storage. a, Principal component analysis of log (CPM + 1)-transformed gene abundances across twelve libraries. Each point represents one sample. Colors identify preservation conditions, and shapes distinguish the two replicates per condition. PC1 and PC2 explain 47.5% and 11.6% of the total variance, respectively. **b,** Pairwise Spearman rank correlations between gene-abundance profiles across a common set of 62,703 genes, retaining zero-abundance values. Cell labels show the correlation coefficients.

Pairwise Spearman correlations across the common set of 62,703 genes provided a complementary measure of agreement in gene-abundance rankings (Fig. 2b and Supplementary Table 2). Correlations between ensilicated and frozen samples ranged from ρ = 0.952 to 0.958, while the two frozen samples correlated at ρ = 0.959. Correlations with frozen samples were lower for bare RNA stored at room temperature (ρ = 0.941–0.946) and at 40 °C (ρ = 0.884–0.927). Within-condition correlations were also higher for ensilicated samples (ρ = 0.956–0.957) than for bare RNA at room temperature (ρ = 0.948) or 40 °C (ρ = 0.914), consistent with the sample comparisons in Supplementary Figs. 3 and 4. These analyses demonstrate closer agreement with frozen gene-level abundance profiles after ensilicated storage compared to bare storage at the same temperatures.

### Ensilicated storage retains annotated transcript information across a broad range of transcript lengths

Gene-level abundance aggregates contributions from multiple transcripts of the same gene and does not capture isoform-level concordance. Annotated-transcript abundance profiles were compared across a common set of 252,913 transcripts (Fig. 3a and Supplementary Table 2). The two frozen replicates exhibited a correlation of ρ = 0.837, while correlations between ensilicated and frozen samples ranged from ρ = 0.793 to 0.837. Correlations with frozen samples were lower for bare RNA stored at room temperature (ρ = 0.744–0.772) and at 40 °C (ρ = 0.414–0.605). Within-condition correlation at 40 °C was also lower for bare RNA (ρ = 0.583) compared to ensilicated RNA (ρ = 0.804). These larger differences at the transcript level indicate that high gene-level correlations do not consistently extend to isoform quantification.

**Figure 3.**
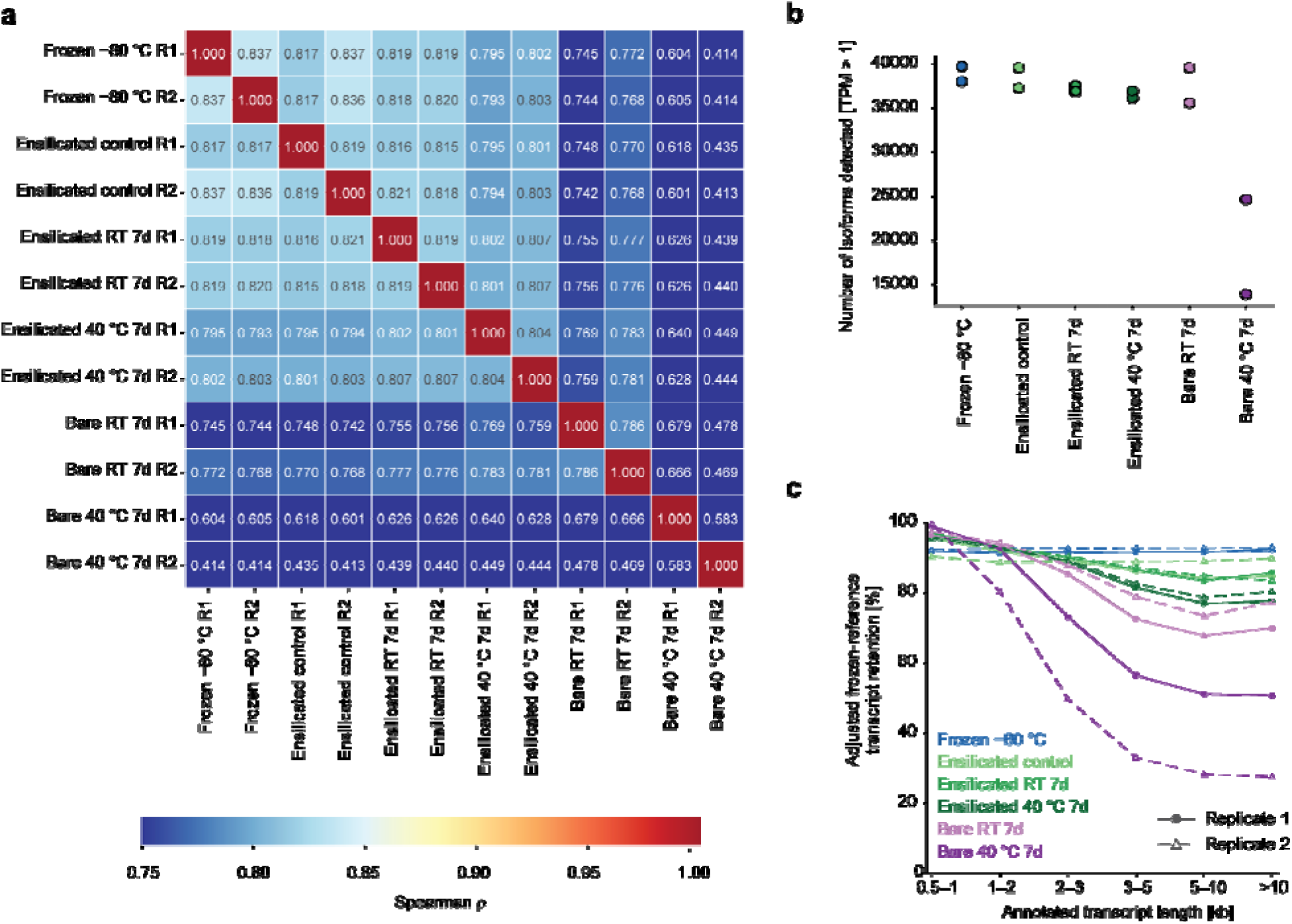
Annotated-transcript abundance, detection, and retention following ensilication and storage. a, Pairwise Spearman rank correlations between annotated-transcript abundance profiles across a common set of 252,913 transcripts, retaining zero-abundance values. Cell labels show correlation coefficients. **b,** Number of annotated transcripts detected at TPM > 1, with two points representing the individual samples per condition. **c,** Expected retention within the fixed frozen-reference set, stratified by annotated exonic length and standardized to a common distribution of frozen starting abundance and annotated isoform count per gene. Retention requires at least one assigned consensus sequence at a common input depth equal to the smallest library. The reference comprises transcripts meeting the detection criteria in both frozen libraries, and transcripts undetected in individual samples remain in the denominator. Circles with solid lines and triangles with dashed lines distinguish the two replicates per condition in **c**.

Frozen and ensilicated samples yielded tens of thousands of detected annotated transcripts, indicating that substantial transcript information remained recoverable from partially degraded RNA samples (Fig. 3b). Bare samples stored at room temperature exhibited overlapping total detection counts, whereas bare samples stored at 40 °C yielded fewer detected transcripts. Subsampling curves demonstrated that gene detection remained high in bare 40 °C samples despite markedly lower transcript detection, highlighting how gene-level summaries can obscure loss of isoform information (Supplementary Fig. 5). However, similar total transcript counts do not confirm recovery of the same transcripts. Retention was therefore assessed against a fixed reference comprising the 13,362 annotated transcripts supported by at least five assignment counts and an abundance of at least 1 CPM in each frozen library. Transcripts undetected after storage remained in this denominator, allowing loss of reference features to contribute directly to the comparison. At a common input depth equal to the smallest consensus-sequence library, expected retention required at least one qualifying sequence per reference transcript (Supplementary Fig. 6a). Frozen samples retained 91.73–92.81% of the reference set, compared with 87.79–89.02% in the ensilicated control. After seven days, ensilicated samples retained 88.17–88.70% at room temperature and 84.55–85.39% at 40 °C. Corresponding retention was lower in bare samples at 78.33–82.90% and 41.95–64.44%, respectively. The comparison depended on the detection threshold. At ten sequences per transcript, one bare 40 °C sample exceeded the ensilicated samples in count-only retention. This reversal was not observed when qualifying sequences were required to support splice-junction combinations that uniquely distinguished the assigned transcript (Supplementary Fig. 6b).

To understand the relationship between transcript length and retention, we used the same frozen-reference population and adjusted for starting abundance and the number of annotated isoforms per gene (Fig. 3c). This comparison retained undetected transcripts within each length stratum and excluded the shortest bin due to the absence of reference transcripts in several adjustment strata. After adjustment, retention in stored samples was lower in the 5–10 kb bin than in the 1–2 kb bin, indicating that the length relationship persisted after accounting for differences in starting representation and annotated isoform complexity. The decline was most pronounced in bare RNA stored at 40 °C. Ensilicated samples retained higher fractions of the longer reference transcripts than bare samples at the same storage temperatures, although retention remained below the frozen reference.

### Gene-abundance differences from frozen are more extensive after bare storage

While correlation and detection analyses provide an overview of agreement, they do not specify which genes exhibit altered abundance. Accordingly, each non-frozen condition was compared with frozen samples using limma moderated t tests on log2(CPM + 1)-transformed gene abundances. Volcano plots were used to summarize the estimated log₂ abundance contrasts and adjusted P values for each comparison (Fig. 4a). Genes were classified as higher or lower in abundance if the estimated log2 contrast exceeded 1 or was less than −1, respectively, and the Benjamini–Hochberg-adjusted P value was below 0.05. Multiple-testing correction was applied across all tested genes and the five contrasts.

**Figure 4.**
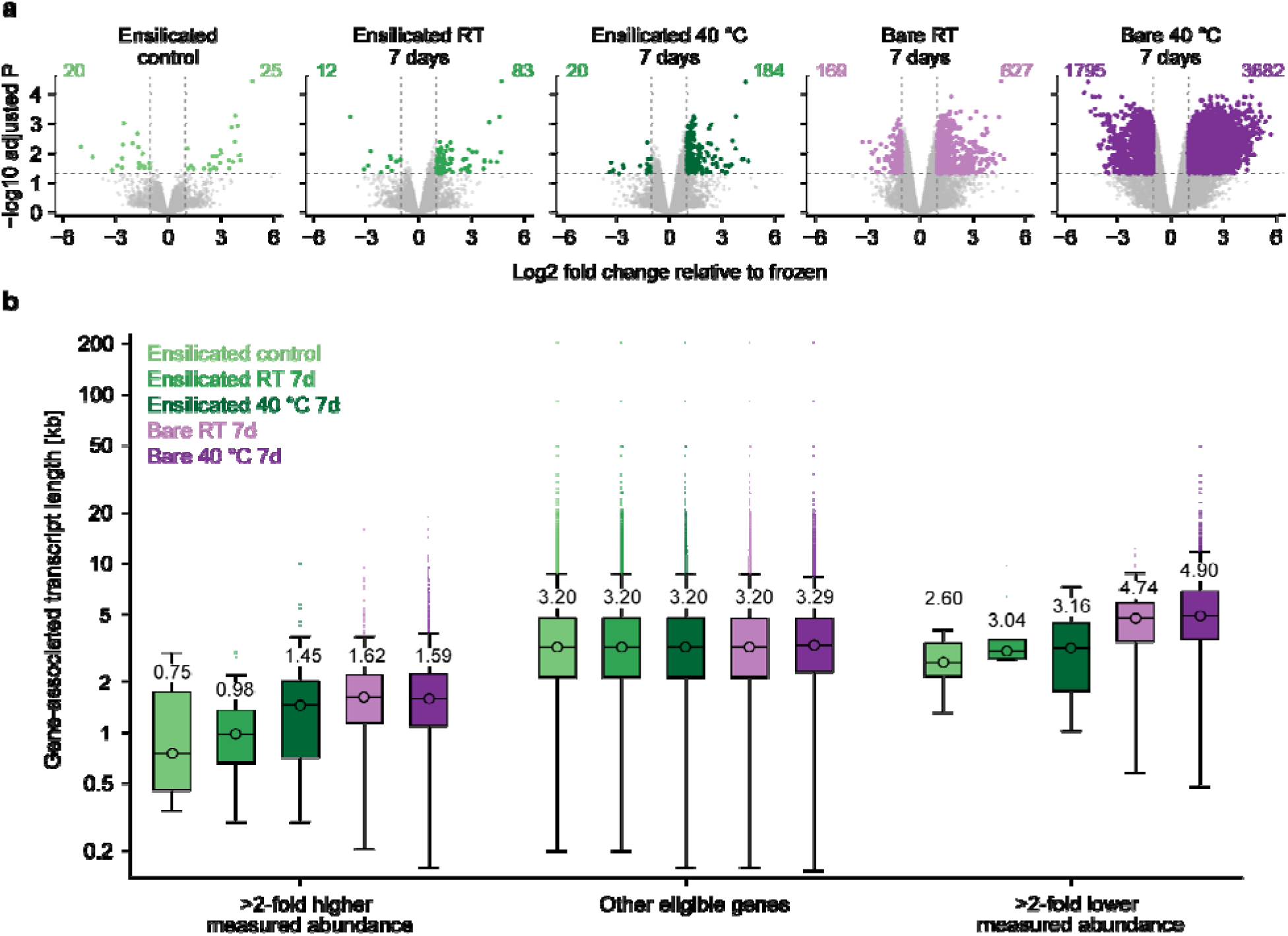
Gene-abundance differences and associated transcript lengths following ensilication and storage. a, Volcano plots compare each non-frozen condition with frozen samples using limma moderated t-tests on log_2_(CPM + 1)-transformed gene abundances. Each point represents a gene with CPM > 5 in at least one library from the displayed condition or frozen reference. The horizontal axis displays the estimated log₂ abundance contrast, and the vertical axis displays −log_10_-adjusted P values. Benjamini–Hochberg correction was applied jointly across all tested genes and five contrasts. Colored points meet both the adjusted P < 0.05 criterion and a log_2_ contrast greater than 1 or less than −1. Remaining genes are shown in grey. Dashed lines indicate these thresholds. Each condition includes two libraries. **b,** Gene-associated transcript-length distributions for the higher-abundance, other eligible, and lower-abundance groups defined in a. Each gene was assigned a fixed exonic length weighted by its annotated isoform abundances in the frozen libraries. Boxes indicate the interquartile ranges, and numerical labels provide median lengths in kilobases.

The ensilicated control exhibited 45 genes meeting both criteria, with 25 showing higher and 20 showing lower measured abundance compared to frozen samples. After seven days of ensilicated storage, these numbers increased to 95 at room temperature (83 higher and 12 lower) and 204 at 40 °C (184 higher and 20 lower). Bare RNA demonstrated substantially more widespread differences, with 796 genes meeting both criteria at room temperature (627 higher and 169 lower) and 5,477 at 40 °C (3,682 higher and 1,795 lower). Although abundance differences were observed following ensilication, fewer genes met the combined statistical and abundance-change criteria than after bare storage at either temperature.

The association between abundance differences and transcript length was subsequently examined (Fig. 4b). Each gene was assigned a fixed exonic length weighted by its annotated isoform abundances in the frozen libraries, ensuring consistent length measures across comparisons. Genes classified as higher in abundance consistently exhibited shorter median lengths in all comparisons, ranging from 0.75 to 1.62 kb, compared to 3.20–3.29 kb among other eligible genes. In bare RNA, genes classified as lower in abundance had median lengths of 4.74 kb after room-temperature storage and 4.90 kb after storage at 40 °C. The corresponding medians in the ensilicated comparisons ranged from 2.60 to 3.16 kb. These findings indicate that abundance differences in bare samples are associated with underrepresentation of genes with longer frozen-reference transcript lengths and relative enrichment of those with shorter lengths.

### Isoform identity and composition after ensilicated storage more closely resemble those observed in frozen samples

Changes in total gene abundance do not fully capture the effects of storage on the relative contributions of isoforms from the same gene. To address this, 2,729 genes with sufficient assignment counts and at least two annotated isoforms reproducibly detected in both frozen libraries were examined (Supplementary Table 2). Each non-frozen sample was compared to the equal-weight mean of the two frozen isoform-proportion vectors. Dominant-isoform discordance ranged from 7.21% to 9.43% in the ensilicated control and from 8.61% to 8.92% after ensilicated storage at room temperature (Fig. 5a). After ensilicated storage at 40 °C, discordance increased to 12.55% and 12.81%, corresponding to 329 of 2,622 and 341 of 2,663 evaluable genes, respectively. Bare samples exhibited higher discordance, ranging from 16.71% to 21.97% at room temperature and from 34.25% to 45.50% at 40 °C. These values corresponded to 793 of 2,315 and 702 of 1,543 evaluable genes, respectively. The direct frozen-pair comparison yielded 6.95% discordance, or 180 of 2,589 genes. Excluding near-tied leading isoforms reduced discordance, particularly in frozen and ensilicated comparisons, whereas bare samples retained higher discordance under both tested margin criteria (Supplementary Fig. 8). These sensitivity analyses differentiate changes involving nearly equal leading isoforms from disagreements that persist when the dominant isoform is more clearly resolved.

**Figure 5.**
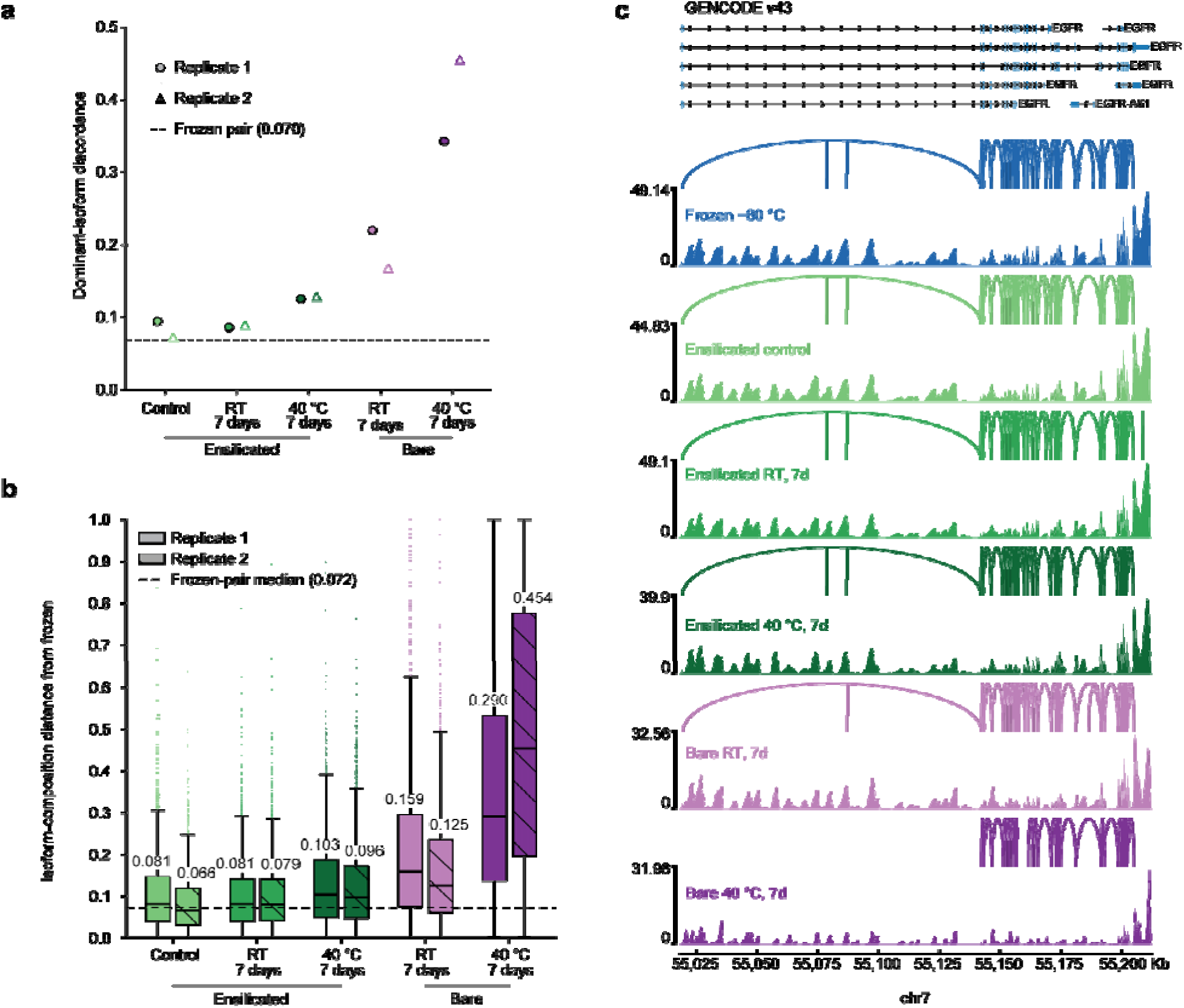
Isoform identity and composition following ensilication and storage. Analyses in **a**,**b** start from 2,729 genes with sufficient assignment counts and reproducible support for multiple annotated isoforms in both frozen libraries. Non-frozen samples are compared with the equal-weight mean of the two frozen isoform-proportion vectors. **a,** Dominant-isoform discordance, defined as the fraction of evaluable genes whose most abundant annotated isoform differs from the frozen reference. Genes absent from the sample or tied for the highest proportion in either compared vector are excluded. Circles and triangles distinguish the two samples per condition. The dashed line indicates the direct frozen-pair discordance (0.070). **b,** Distributions of within-gene isoform-composition distance, ranging from zero for identical proportions to one for nonoverlapping composition. Boxes show interquartile ranges, and horizontal lines and numerical labels indicate medians. Solid and hatched boxes distinguish the two samples per condition. The dashed line indicates the direct frozen-pair median (0.072). Genes without assigned transcript counts have undefined distances and are reported separately. Sample-specific denominators are provided in Supplementary Table 2 and Supplementary Fig. 8. **c,** Genome browser view of the EGFR locus with GENCODE v43 annotation. Filled tracks show CPM-normalised coverage averaged across the two samples per condition, with track-specific scales indicated. Arcs summarise support for individual splice junctions.

The magnitude of redistribution across within-gene isoform proportions was quantified using composition distance, defined as half the sum of the absolute differences between sample and reference proportions (Fig. 5b). A value of zero indicates identical proportions, while a value of one indicates nonoverlapping composition. The direct frozen-pair median composition distance was 0.072, with an interquartile interval of 0.035 to 0.132. In the ensilicated control, median distances and interquartile intervals were 0.081 (0.040 to 0.147) and 0.066 (0.032 to 0.118). After ensilicated storage, the corresponding values were 0.081 (0.040 to 0.141) and 0.079 (0.042 to 0.140) at room temperature, and 0.103 (0.049 to 0.186) and 0.096 (0.047 to 0.172) at 40 °C. Bare RNA exhibited larger composition shifts, with values of 0.159 (0.074 to 0.296) and 0.125 (0.061 to 0.234) at room temperature, and 0.290 (0.137 to 0.533) and 0.454 (0.196 to 0.776) at 40 °C. These distributions represent genes with assigned transcript counts. The larger shifts observed in bare RNA were accompanied by a loss of evaluable genes, with 28 and 6 reference-eligible genes lacking assigned counts at room temperature and 185 and 1,006 at 40 °C, compared with 0 to 4 per ensilicated sample.

### Ensilicated samples retain a greater number of annotated transcripts with strict structural support

To assess how differences in transcript representation correspond to coverage of annotated transcript ends, the median missing annotated exonic sequence at the 5′ end was measured among uniquely assigned consensus sequences with evaluable boundaries (Fig. 6a). Frozen samples exhibited median missing values of 0.899 and 0.907 kb at the 5′ end. In comparison, medians ranged from 0.934 to 0.988 kb in ensilicated conditions and from 0.945 to 0.994 kb in bare samples stored at room temperature. Bare samples stored at 40 °C demonstrated greater missing 5′ sequence, with medians of 1.319 and 1.535 kb. This increase was associated with smaller amounts of missing annotated sequence at the 3′ end, with medians of 0.078 and 0.007 kb, compared to 0.244 and 0.229 kb in frozen samples. The observed pattern of reduced upstream coverage and closer agreement with annotated 3′ ends is consistent with oligo-dT capture of poly(A)-bearing fragments after fragmentation. However, these results do not establish the direction of molecular cleavage.

**Figure 6.**
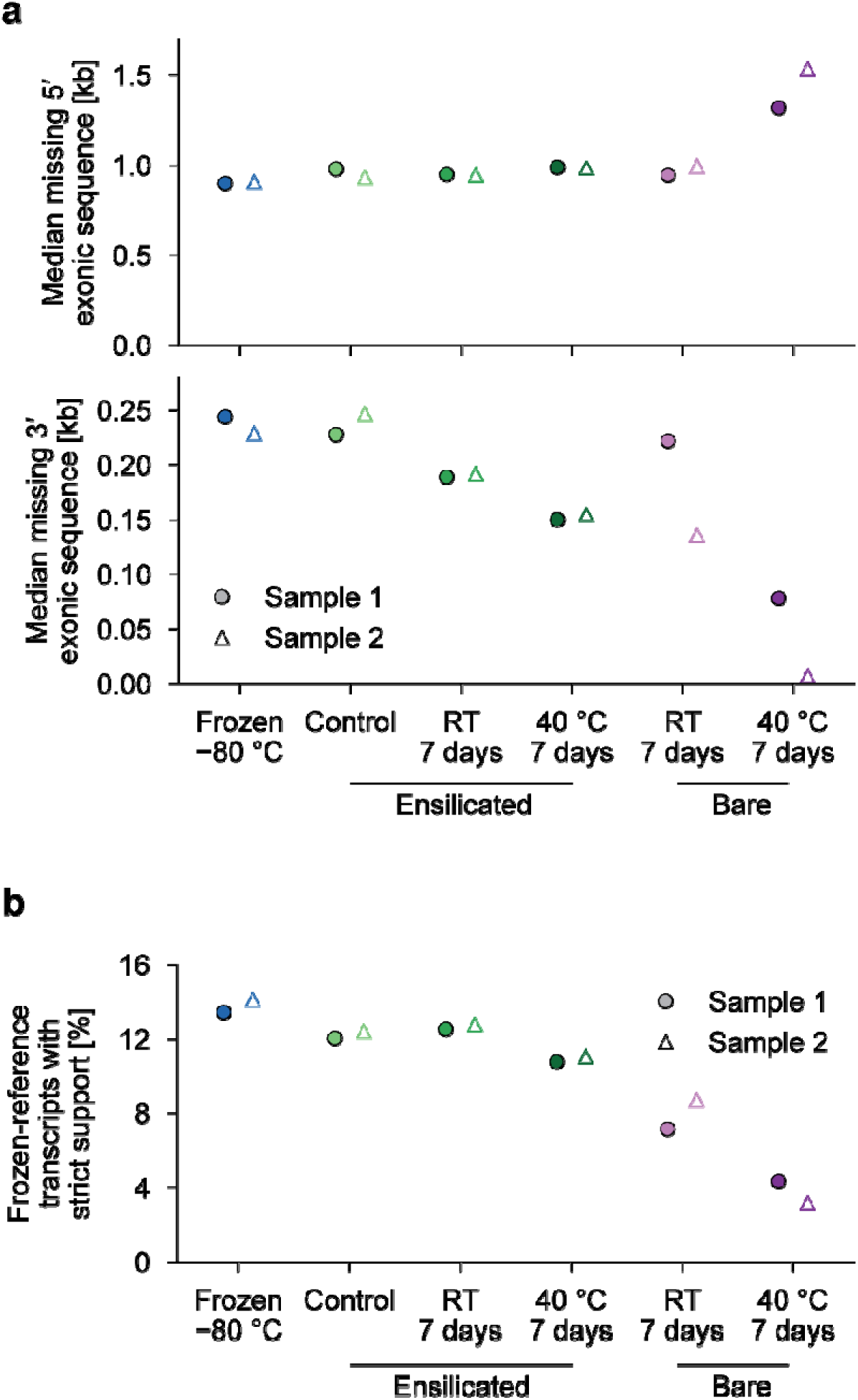
Annotated transcript-end coverage and structural support following ensilication and storage. a, Median missing annotated exonic sequence at the 5′ end (upper panel) and 3′ end (lower panel), measured among uniquely assigned consensus sequences with evaluable alignment and annotation boundaries. Missing lengths represent annotated exonic bases beyond the corresponding alignment end, accounting for transcript strand and excluding intronic sequence. **b,** Expected percentage of frozen-reference transcripts supported by at least one consensus sequence meeting the strict structural criteria. Qualifying sequences must support a complete, uniquely distinguishing annotated intron chain and have both original genomic alignment ends within 50 nucleotides of the annotated boundaries. Retention is evaluated at a common input depth equal to the smallest consensus-sequence library. All samples use the same fixed reference set of transcripts reproducibly detected in both frozen libraries, including transcripts without qualifying support in individual samples. Filled circles and open triangles distinguish replicate 1 and 2 within each condition.

To evaluate whether the preservation comparison remained consistent when transcript detection required stronger structural evidence, qualifying sequences were restricted to those supporting a complete, uniquely distinguishing annotated intron chain and both genomic alignment ends within 50 nucleotides of the annotated boundaries (Fig. 6b). Expected retention was defined as the presence of at least one qualifying consensus sequence at the same common input depth used for the primary retention analysis. All samples used the same frozen-reference denominator, including transcripts without qualifying support. Under these criteria, frozen samples retained 13.44–14.13% of reference transcripts, compared to 12.05–12.43% in the ensilicated control. After seven days of ensilicated storage, strict retention was 12.53–12.79% at room temperature and 10.77–11.05% at 40 °C. Bare samples retained lower fractions at both temperatures, with 7.16–8.72% at room temperature and 3.19–4.33% at 40 °C. These findings indicate that greater retention in ensilicated samples persisted when recovery required evidence for both distinguishing splice structure and annotated transcript boundaries.

Imposing requirements for complete intron chains and agreement with annotated transcript boundaries reduced the proportion of qualifying consensus sequences across all conditions (Supplementary Fig. 7). At the observed library depth, 0.88–0.92% of frozen input consensus sequences met the strict criteria, compared to 0.41–0.51% in bare samples stored at 40 °C. These percentages represent the fraction of input sequences meeting the structural criteria, while Figure 6b reports the fraction of frozen-reference transcripts supported by such sequences. The presence of missing annotated end sequence in frozen samples further demonstrates that pipeline-defined FLNC status does not guarantee complete annotated-transcript coverage. These measurements quantify the support for annotated transcript features among captured sequences.

## Discussion

Ensilicated samples retained a greater amount of recoverable annotated transcript information than bare RNA after seven days at either room temperature or 40 °C. Long-read sequencing demonstrated this protective effect by revealing differences in transcript retention, isoform composition, and the structural features that support transcript identification. These measurements expand upon previous approaches for evaluating ambient RNA preservation. Prior studies of RNAlater-preserved tissues have utilized microarrays and targeted RNA quality assessments [11, 12], while dry-storage matrices have been evaluated using integrity measurements, RT-qPCR, microarrays, and short-read RNA sequencing [10, 15, 16]. By analyzing transcriptome-wide isoform representation in conjunction with sequence-level structural support, the present study assesses preservation aspects that are not fully captured by bulk integrity or gene-level summaries.

The significance of this additional information was evident when gene-level agreement was evaluated alongside isoform recovery. Gene-level correlations remained relatively high even in bare samples that exhibited substantial transcript loss and altered isoform composition. Although captured sequences may still identify a gene, they may lack the necessary features to distinguish between its isoforms. Furthermore, degradation can distort relative gene abundance, as previously demonstrated in short-read RNA sequencing [6]. In this study, 5,477 genes met the statistical significance and abundance-change criteria in bare RNA at 40 °C, compared to 204 genes in ensilicated RNA under the same conditions. Since the samples originated from a shared RNA source, these differences and the associated changes in isoform composition reflect altered measurement of the starting material. Reporting absent genes and analyzing near-tied dominant isoforms further differentiated information loss and substantial redistribution from minor changes in relative abundance.

Interpreting this loss required a reference that remained constant across storage conditions. Counting all detected transcripts conflates retention of previously measurable features with differences in which transcripts surpass the detection threshold. The frozen-reference analysis instead included missing transcripts in the denominator and demonstrated lower retention of longer transcripts after adjusting for starting abundance and annotated isoform complexity. This approach resolved the apparent advantage for long transcripts observed with the original annotation-wide denominator. However, detection remained sensitive to the chosen counting criterion. At the ten-sequence threshold, one bare 40 °C sample exhibited slightly higher count-only retention than the ensilicated samples, but this difference was eliminated when qualifying sequences were required to support distinguishing splice junctions (Supplementary Fig. 6). Therefore, a higher count for an assigned transcript did not necessarily provide stronger evidence for its isoform structure.

The structural analyses directly addressed this distinction. Greater retention in ensilicated samples persisted when qualifying sequences were required to support a complete, uniquely distinguishing intron chain and to align with both annotated transcript boundaries. This criterion provides stronger evidence of recovered transcript structure than pipeline-defined FLNC status alone. Because a poly(A)-bearing RNA fragment can generate cDNA carrying both library primers, FLNC classification does not confirm preservation of both native transcript ends. In line with this distinction, bare 40 °C samples exhibited increased loss of annotated sequence at the 5′ end, consistent with preferential capture of poly(A)-bearing fragments without indicating the direction of molecular cleavage. Similarly, a nanopore direct RNA sequencing degradation study reported reduced library complexity and overrepresentation of shorter genes and isoforms [9]. However, incomplete annotated coverage was also observed in the frozen samples, highlighting the importance of the reference material’s condition in interpreting what was protected.

Frozen samples exhibited RINe values of 4.9–5.0 yet retained substantial gene and annotated-transcript information, including transcripts supported by complete intron chains and annotated boundaries. Preventing further loss of this information remains a meaningful preservation objective, even when the starting material is partially degraded. The strict structural criteria selected only a small subset of captured sequences, in part because monoexon transcripts and isoforms unresolved by junction evidence were excluded. Consequently, the pass fractions cannot be interpreted as the proportion of original RNA molecules that remain intact. Collectively, these observations define the study as a comparison of recoverable information in the tested preparations, rather than establishing performance with initially high-integrity RNA or equivalence to frozen storage.

The strength of this comparison is also influenced by the experimental design. A single pooled RNA stock served as a common source, and all twelve libraries were prepared simultaneously, with the ensilicated control and stored ensilicated samples released on the same day. However, replication was limited to two samples per condition, and each condition was sequenced in a separate pool, which precluded separation of condition and pool effects. The abundance tests assume independent library errors, while depth matching addresses sequencing-depth differences without increasing experimental replication. As a result, the observed differences reflect the combined effects of preservation and sequencing, rather than establishing absolute RNA recovery or distinguishing physical RNA loss from capture, clustering, and assignment effects.

Within this scope, the principal finding is that ensilicated, purified RNA retained more transcript information than bare RNA over the seven-day storage interval. The room-temperature comparison directly supports this interval, while the 40 °C condition assesses elevated-temperature stress without establishing longer-term shelf life. Implementing this storage benefit in a complete workflow also requires consideration of the preparation process. The evaluated protocol involved two days of mixing at 1,400 rpm followed by subsequent release, which limits immediate application in settings lacking powered equipment. Compatibility with intact tissues and crude lysates remains unresolved. The demonstrated contribution is therefore protection during storage after preparation, as supported by long-read measurements that link the amount of transcript information recovered to the structural evidence underlying its identification.

## Conclusion

Long-read RNA sequencing demonstrated that ensilication preserves recoverable transcript information during storage. After seven days at room temperature or 40 °C, ensilicated samples retained a higher number of annotated transcripts and exhibited smaller deviations from frozen-reference gene abundance and isoform composition compared to untreated RNA. Enhanced transcript retention was also observed when qualifying sequences were required to support a complete, uniquely distinguishing intron chain and to match both annotated transcript boundaries. Collectively, these results indicate that ensilication effectively limits further loss of transcript information during storage of partially degraded RNA at ambient temperature. By integrating transcriptome-wide quantification with structural analysis, this study highlights the utility of long-read sequencing for assessing RNA preservation beyond bulk integrity and gene-level detection.

## Declarations

### Ethics approval and consent to participate

The RNA used in this study was obtained commercially (BioChain Universal Human RNA) as a deidentified, pooled reagent derived from human tissue. The study did not involve the collection of samples or data from identifiable living individuals and did not constitute human subjects research; ethics approval was therefore not required.

## Consent for publication

Not applicable.

## Availability of data and materials

The datasets generated and analysed during the current study are available in the NCBI Sequence Read Archive under BioProject accession PRJNA1447563 (https://www.ncbi.nlm.nih.gov/bioproject/PRJNA1447563). All analysis scripts and computational workflows used to generate the figures and results are available at https://github.com/jearlbcache/ensilication-rna. Ensilication and de-ensilication reagents are available from Cache DNA, Inc. on request.

## Competing interests

J.L.B. holds equity in and is a co-founder of Cache DNA, Inc.; consults for OpenAI and Glyphic Biotechnologies. M.M. and H.G. are employees of Signios Biosciences.

## Funding

J.L.B. received support from PacBio in the form of a sequencing grant.

## Authors’ contributions

M.M. performed sample preparation and sequencing. H.G. supervised sample preparation and sequencing. J.L.B. performed the bioinformatic analyses, conceived the study, and wrote the manuscript. All authors read and approved the final manuscript.

## Acknowledgements

The authors thank PacBio for support through a sequencing grant.

## Authors’ information

Not applicable.

## Notes

### Competing Interest Statement

J.L.B. holds equity and is the co-founder of Cache DNA, Inc.; consults for OpenAI and Glyphic Biotechnologies. M.M. and H.G. are employees of Signios Biosciences.

### Summary of Updates

Title, abstract and main text revised to focus on transcriptome-wide information preservation; starting RNA integrity and the scope of conclusions clarified; Methods expanded to describe sample handling, sequencing, normalization and statistical analyses; Figure 1c updated with individual sample medians and interquartile intervals; Figure 3c revised to assess frozen-reference transcript retention with adjustment for starting abundance and isoform complexity; Figure 4 updated with statistical testing of gene-abundance differences and associated transcript-length distributions; Figures 5-6 revised to quantify isoform-composition changes, annotated transcript-end coverage and structural support; supplementary information updated with detection-threshold, sequencing-depth, structural-support and dominant-isoform sensitivity analyses, together with analysis denominators; figure captions, software citations and author affiliations updated.

