## Supplementary Information for "Evaluating transcriptome-wide information preservation by ensilication using long-read RNA sequencing"

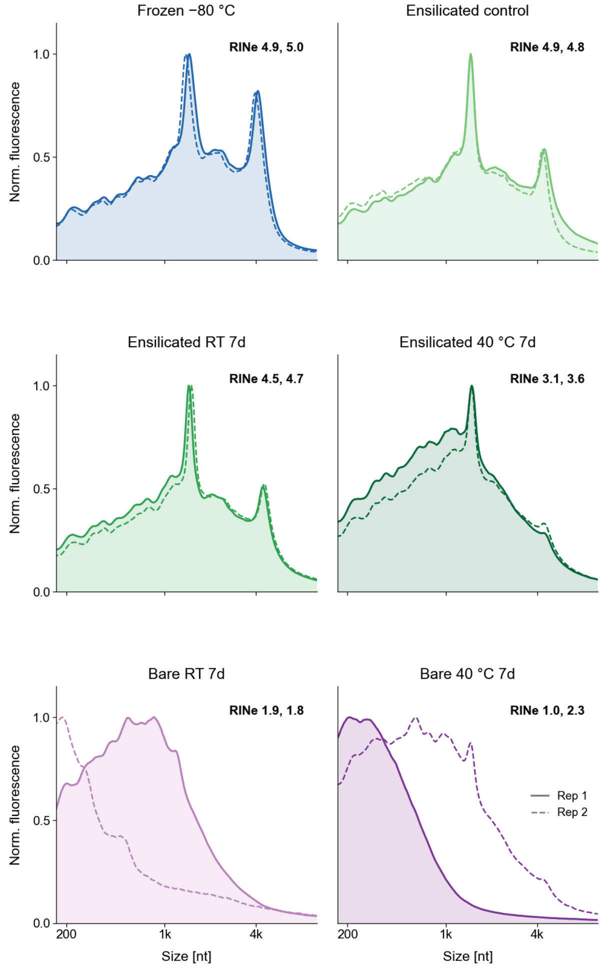


**Supplementary Figure 1. TapeStation electropherograms for all 12 samples.**

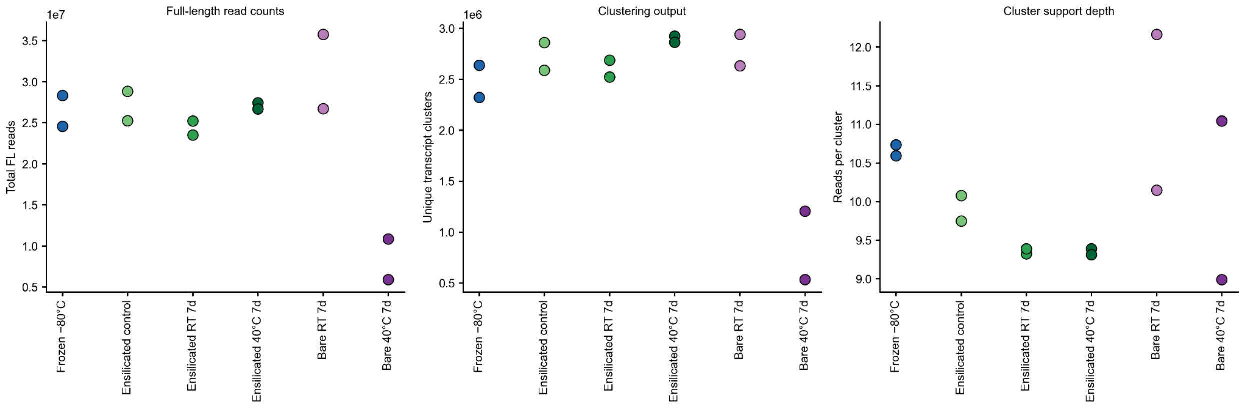


**Supplementary Figure 2. Sequencing yield and clustering summary across samples.** Numbers of full-length non-chimeric (FLNC) reads (left), consensus sequences produced by clustering (middle), and FLNC reads per cluster (right). Each point represents one of two samples per preservation condition.

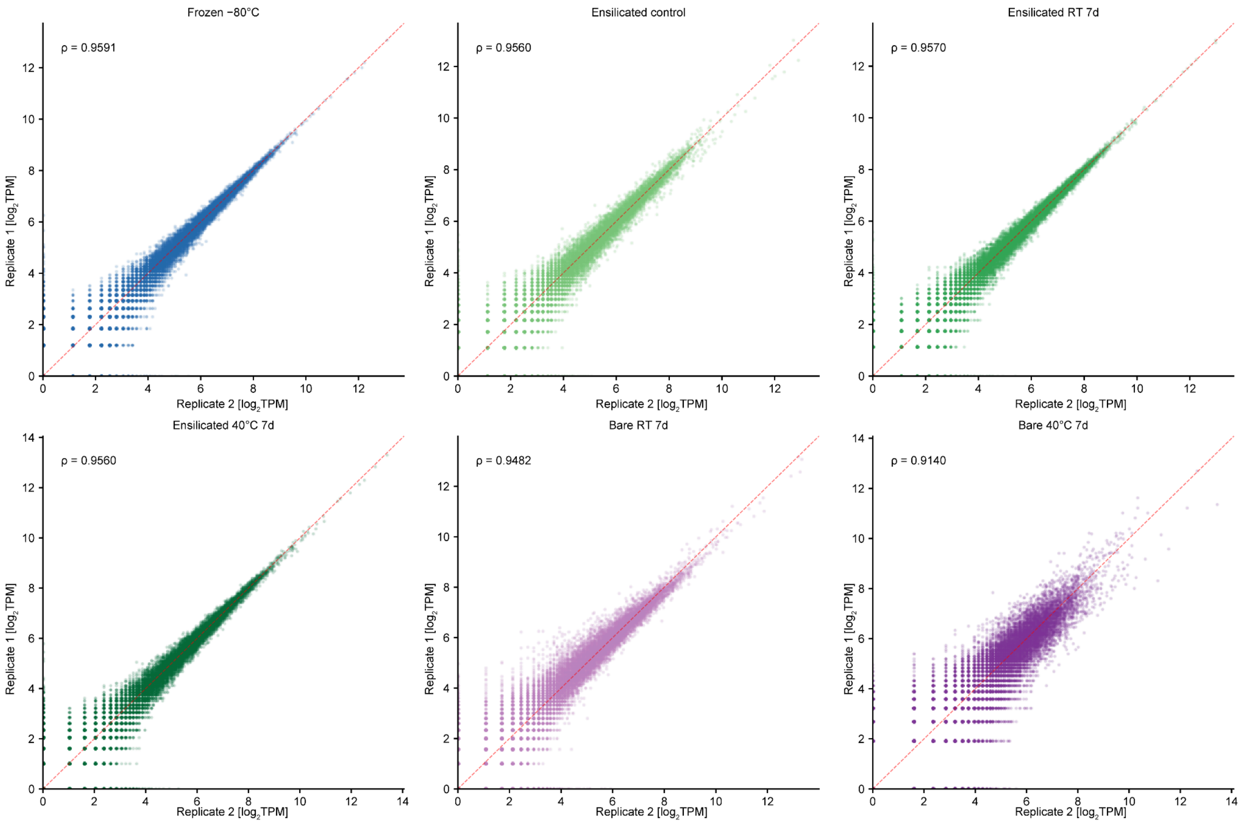


**Supplementary Figure 3. Replicate concordance.** Pairwise scatter plots of gene-level $log_2$TPM between replicates within each condition.

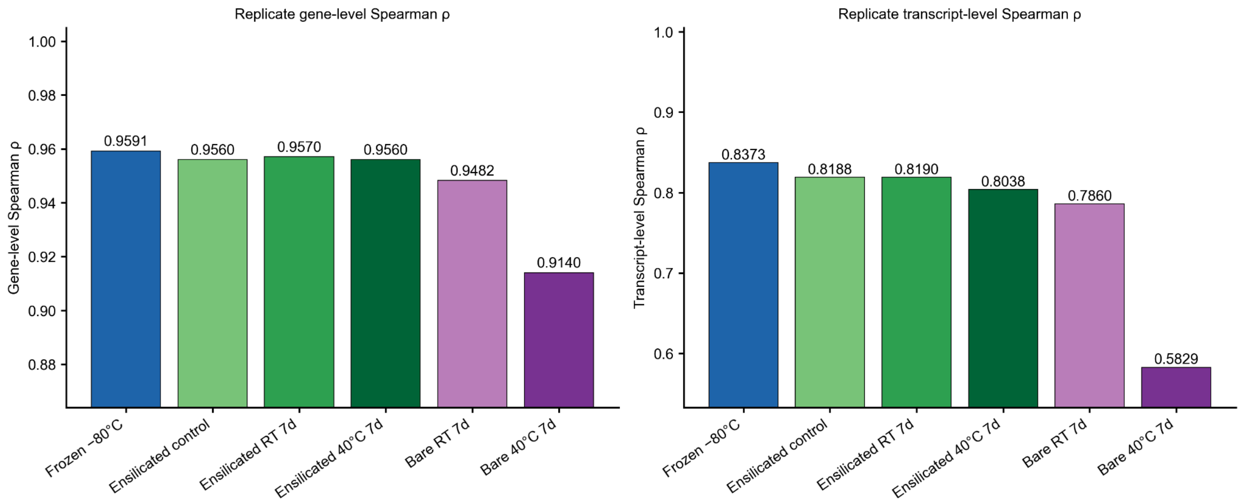


**Supplementary Figure 4. Replicate concordance.** Within-condition replicate Spearman correlation at the gene level (left) and transcript level (right) for all six conditions.

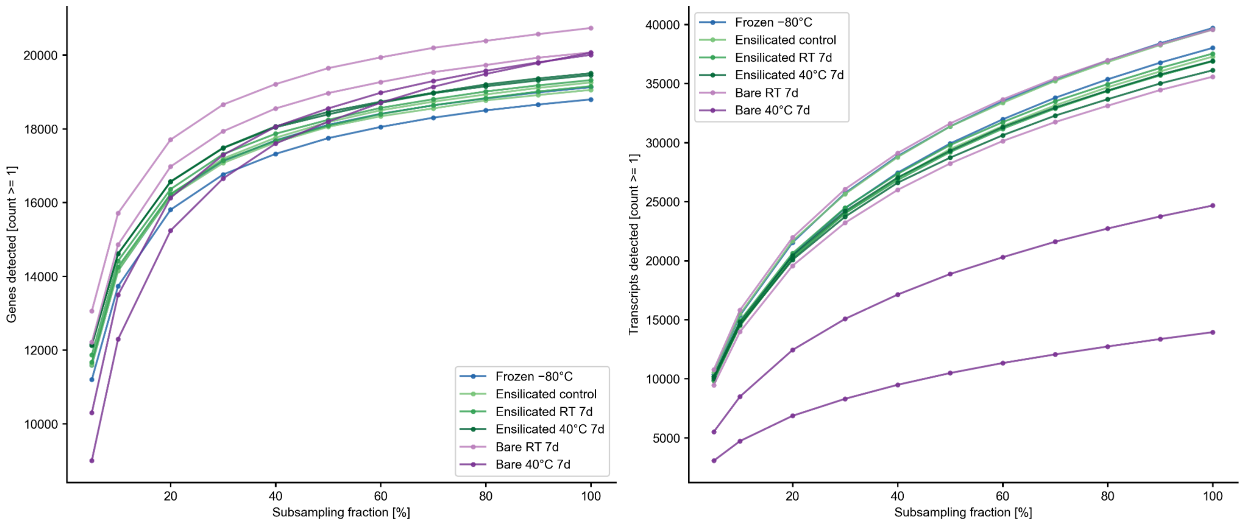


**Supplementary Figure 5. Gene and transcript detection saturation.** Number of genes (left) and transcripts (right) detected as a function of subsampled read depth for each condition.

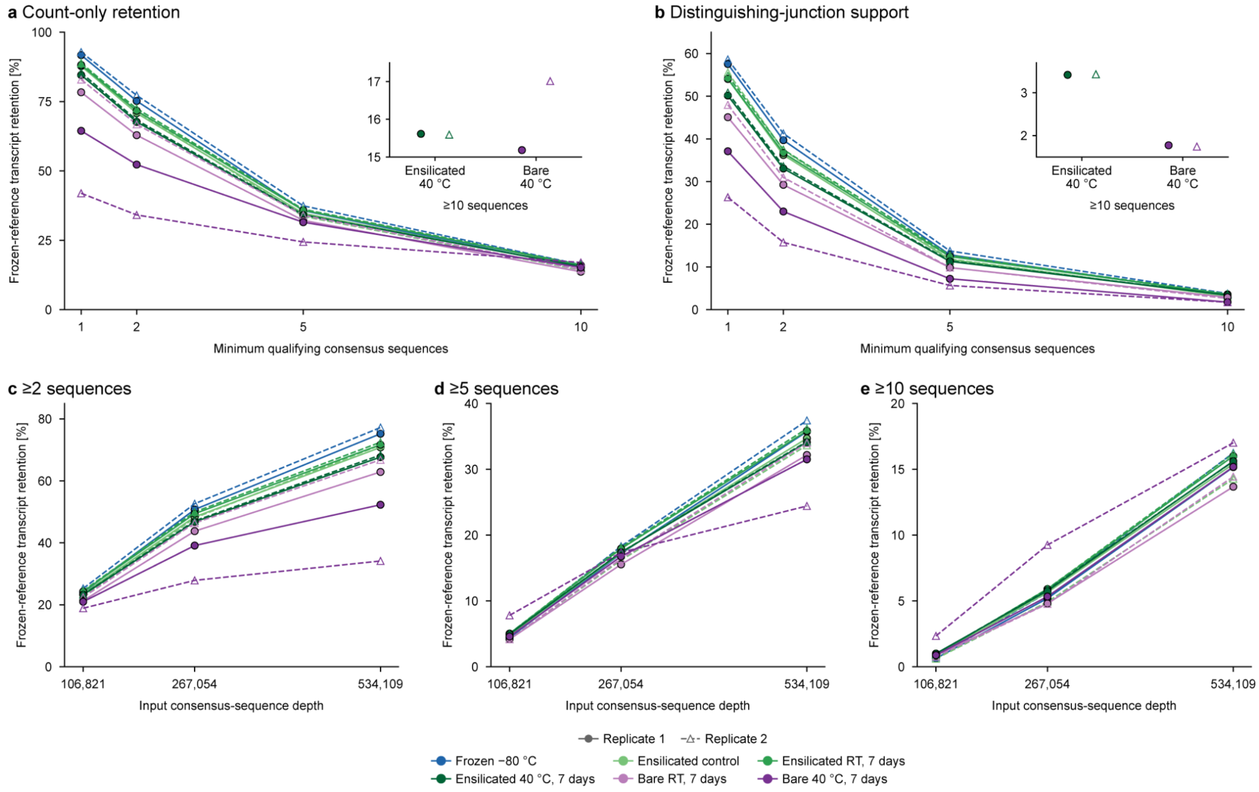


**Supplementary Figure 6. Sensitivity of annotated transcript retention to detection criteria and sequencing depth.** A fixed reference set was defined from annotated transcripts reproducibly detected in both frozen libraries, yielding 13,362 transcripts. All reference transcripts remain in the denominator, including those undetected in individual samples. Retention is the expected number meeting the detection criterion, expressed as a percentage of this reference set. **a,b,** Expected retention after standardizing input depth to the number of consensus sequences available in the smallest library (534,109), assessed across detection thresholds requiring at least 1, 2, 5 or 10 qualifying sequences per transcript. **a,** Count-only retention based on the original IsoQuant annotated-transcript counts. **b,** Retention requiring observed splice-junction combinations consistent with, and uniquely distinguishing, the assigned annotated transcript. Insets show the ensilicated and bare 40 °C comparisons at the ten-sequence threshold. **c–e,** Count-only retention at input depths corresponding to approximately 20%, 50% and 100% of the common depth, shown separately for detection thresholds of at least two (**c**), five (**d**) or ten (**e**) sequences per transcript. Filled circles with solid lines and open triangles with dashed lines distinguish the two samples per condition. Lines connect estimates for the same sample across the evaluated thresholds or depths.

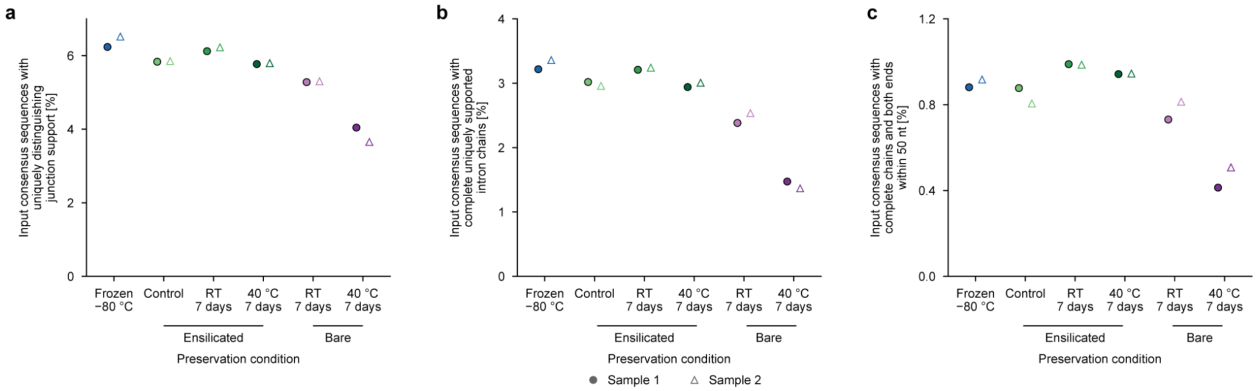


**Supplementary Figure 7. Structural support for annotated transcript assignments.** Percentages of input consensus sequences satisfying progressively stricter structural-support criteria. **a,** Sequences whose observed splice-junction combinations are consistent with, and uniquely distinguish, the assigned annotated transcript. **b,** Sequences additionally supporting the complete annotated intron chain. **c,** Sequences satisfying the complete-chain criterion with both genomic alignment ends within 50 nucleotides of the corresponding annotated transcript boundaries. Within each sample, all three panels use the total number of input consensus sequences at full observed sequencing depth as the denominator, including sequences that fail the support criteria. Each consensus sequence contributes once, without weighting by cluster-member counts. Each point represents one of two samples per condition. Filled circles indicate Replicate 1 and open triangles indicate Replicate 2.

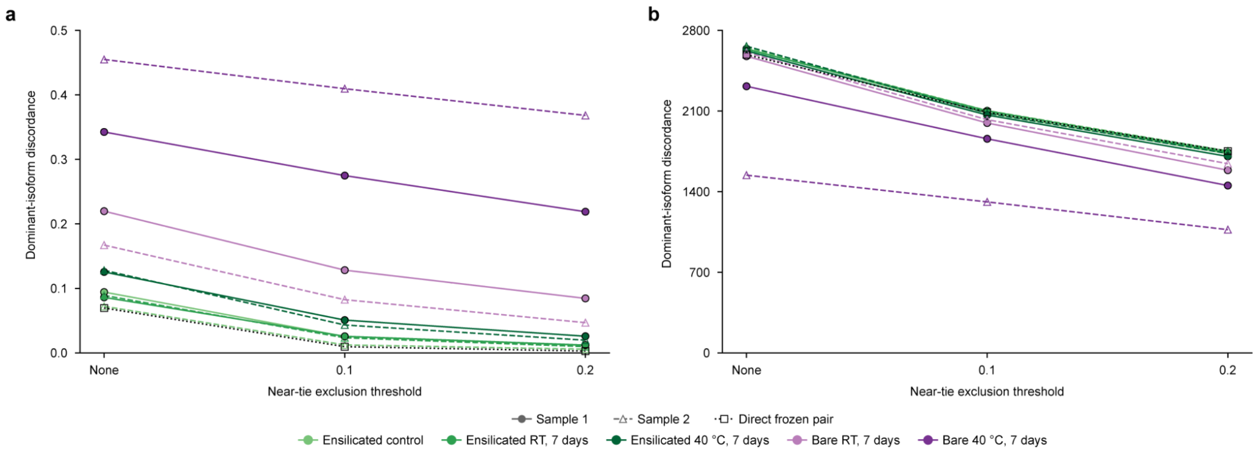


**Supplementary Figure 8. Sensitivity of dominant-isoform discordance to near-tie exclusion.** To restrict comparisons to genes with reproducible multi-isoform support, eligibility required at least 20 annotated-transcript assignment counts in each frozen library and at least two annotated isoforms each supported by two or more counts in both frozen libraries. These criteria yielded 2,729 genes before sample-specific exclusions. **a,** Fraction of evaluable genes whose highest-proportion annotated isoform differs between each sample and the frozen reference, defined by the equal-weight mean of the two frozen samples' within-gene isoform proportions. **b,** Corresponding numbers of evaluable genes. Near-tie exclusion tests whether discordance persists when the leading isoform clearly exceeds the second-ranked isoform. Thresholds of 0.1 and 0.2 require this difference to exceed 10 and 20 percentage points, respectively, in both the sample and the frozen reference. "None" indicates no additional near-tie exclusion. Genes with tied leading isoforms or zero assigned counts are excluded throughout. The dotted line with open squares compares the two frozen samples directly, using their own eligible gene subset under the same exclusion rule at each threshold. Filled circles with solid lines and open triangles with dashed lines distinguish the two samples per condition.

**Supplementary Table 1. RNA recovery**. Samples were measured by Nanodrop

| Preservation | Condition | Replicate | Recovery [%] | A260/A280 | A260/A230 |
|---|---|---|---|---|---|
| Frozen | –80 °C | 1 | 114.4 | 2.092 | 2.177 |
| Frozen | –80 °C | 2 | 174.7 | 2.075 | 2.177 |
| Ensilicated | Control | 1 | 93.5 | 2.052 | 1.586 |
| Ensilicated | Control | 2 | 91.4 | 2.065 | 1.552 |
| Ensilicated | RT 7d | 1 | 105.1 | 2.054 | 1.494 |
| Ensilicated | RT 7d | 2 | 98.8 | 2.035 | 1.469 |
| Ensilicated | 40 °C 7d | 1 | 107.7 | 2.025 | 2.144 |
| Ensilicated | 40 °C 7d | 2 | 98.2 | 2.037 | 2.145 |
| Bare | 40 °C 7d | 1 | 6.1 | 2.105 | 2.554 |
| Bare | 40 °C 7d | 2 | 1.8 | 1.32 | 1.711 |
| Bare | RT 7d | 1 | 41.3 | 2.032 | 1.685 |
| Bare | RT 7d | 2 | 49.3 | 2.06 | 1.824 |

**Supplementary Table 2. Numbers of genes and transcripts included in correlation and isoform-composition analyses.**

| Figure/panel | Analysis | Sample or comparison | Feature type | Number included |
|---|---|---|---|---|
| Figure 2b | Gene-level Spearman correlation | Every pairwise comparison among 12 libraries | genes | 62,703 |
| S3 and S4 left | Within-condition gene correlations | Every within-condition Sample 1 vs Sample 2 comparison | genes | 62,703 |
| Figure 3a | Transcript-level Spearman correlation | Every pairwise comparison among 12 libraries | transcripts | 252,913 |
| S4 right | Within-condition transcript correlations | Every within-condition Sample 1 vs Sample 2 comparison | transcripts | 252,913 |
| Figure 5b | Isoform-composition distance | Ensilicated control, Sample 1 | Multi-isoform genes | 2,727 |
| Figure 5b | Isoform-composition distance | Ensilicated control, Sample 2 | Multi-isoform genes | 2,729 |
| Figure 5b | Isoform-composition distance | Ensilicated RT, 7 days, Sample 1 | Multi-isoform genes | 2,729 |
| Figure 5b | Isoform-composition distance | Ensilicated RT, 7 days, Sample 2 | Multi-isoform genes | 2,728 |
| Figure 5b | Isoform-composition distance | Ensilicated 40 °C, 7 days, Sample 1 | Multi-isoform genes | 2,725 |
| Figure 5b | Isoform-composition distance | Ensilicated 40 °C, 7 days, Sample 2 | Multi-isoform genes | 2,728 |
| Figure 5b | Isoform-composition distance | Bare RT, 7 days, Sample 1 | Multi-isoform genes | 2,701 |
| Figure 5b | Isoform-composition distance | Bare RT, 7 days, Sample 2 | Multi-isoform genes | 2,723 |
| Figure 5b | Isoform-composition distance | Bare 40 °C, 7 days, Sample 1 | Multi-isoform genes | 2,544 |
| Figure 5b | Isoform-composition distance | Bare 40 °C, 7 days, Sample 2 | Multi-isoform genes | 1,723 |
| Figure 5b | Isoform-composition distance | Frozen Sample 2 vs Sample 1 | Multi-isoform genes | 2,729 |

1. The frozen-eligible multi-isoform set contains 2,729 genes before sample-specific exclusions. Each gene has at least 20 annotated-transcript assignment counts in each frozen library and at least two identical transcript IDs with at least 2 counts in both. Genes with zero sample-assigned counts have undefined composition and are excluded from Figure 5b. Non-frozen samples compare with the equal-weight mean frozen proportion vector.

2. Counts refer to genuine genes or transcripts. Figure 2b excludes the all-zero __unassigned summary row. Existing correlations in Figure 3a, S3 and S4 retain this additional non-feature row, giving 62,704 gene-level or 252,914 transcript-level input rows. The summary row does not contribute to the feature counts reported here.
